# Illuminating spatial A-to-I RNA editing signatures within the Drosophila brain

**DOI:** 10.1101/370296

**Authors:** Anne L. Sapiro, Anat Shmueli, Gilbert Lee Henry, Qin Li, Tali Shalit, Orly Yaron, Yoav Paas, Jin Billy Li, Galit Shohat-Ophir

**Author notes:** Correspondence should be addressed to G.S.O or to J.B.L. Authors contributed equally to this work (Anne L. Sapiro, Anat Shmueli). Current Address, Cold Spring Harbor Laboratory New York, USA.

## Abstract

**Summary:** Adenosine-to-inosine (A-to-I) RNA editing, catalyzed by ADAR enzymes, is a ubiquitous mechanism that generates transcriptomic diversity. This process is particularly important for proper neuronal function; however, little is known about how RNA editing is dynamically regulated between the many functionally distinct neuronal populations of the brain. In this resource paper, we present a spatial RNA editing map in the *Drosophila* brain and show that different neuronal populations possess distinct RNA editing signatures. After purifying and sequencing RNA from genetically marked neuronal nuclei, we identified a large number of novel editing sites and compared editing levels in hundreds of transcripts across nine functionally different neuronal populations. We found distinct editing repertoires for each population, including novel sites in repeat regions of the transcriptome and differential editing in highly conserved and likely functional regions of transcripts that encode essential neuronal genes. These changes are site-specific and not driven by changes in *Adar* expression, suggesting a complex, targeted regulation of editing levels in key transcripts. This fine-tuning of the transcriptome between different neurons by RNA editing may account for functional differences between distinct populations in the brain.

**Significance Statement:** A fundamental question in contemporary neuroscience is how the remarkable cellular diversity required for the intricate function of the nervous system is achieved. In this manuscript, we bridge the gap between a cellular machinery that is known to diversify the transcriptome and the existence of distinct neuronal populations that compose *Drosophila* brain. Adenosine-to-inosine (A-to-I) RNA-editing is a ubiquitous mechanism that generates transcriptomic diversity in cells by recoding certain adenosines within the pre-mRNA sequence into inosines. We present a spatial map of RNA editing across different neuronal populations in *Drosophila* brain. Each neuronal population has a distinct editing signature, with the majority of differential editing occurring in highly conserved regions of transcripts that encode ion channels and other essential neuronal genes.

## Introduction

The complexity and function of the nervous system is due in part to the existence of various types of neuronal cells with distinct functions, anatomical locations, structures, physiologies, and connectivity. This diversity is accomplished by molecular programs that shape the repertoire of RNA molecules and proteins within each cell, giving rise to populations with distinct molecular signatures. Numerous mechanisms can contribute to the proteomic diversity between neuronal populations, including RNA modifications. One particular modification that is critical to brain function is adenosine-to-inosine (A-to-I) RNA editing, catalyzed by a class of proteins called Adenosine Deaminases that act on RNA (ADARs), which are conserved across metazoans^1,2^. The resulting inosines are read by the cellular machinery as guanosines, leading to a variety of consequences including altered splicing and gene expression and changes to the amino acid sequences of proteins^3,4^.

Thousands of RNA editing sites have recently been discovered in *Drosophila*^5-12^, and the loss of ADAR editing results in mainly neuronal and behavioral phenotypes^2,13^. Many of these sites are predicted to cause nonsynonymous protein coding (“recoding”) changes in genes that are expressed and function primarily in neurons, such as ion channels and pre-synaptic proteins involved in neurotransmission. Evolutionary analysis of editing levels across multiple *Drosophila* species indicates that many of these protein coding changes in neuronal genes are being selected for over evolution, suggesting their editing may be functionally important^9-11^.

Studies indicate that editing modulates the kinetics of the voltage-dependent K+ channels *Shaker* and *Shab*^14,15^, the agonist potency of the GABA-gated Cl^-^ channel, *Rdl* ^16^, and the voltage sensitivity and closing kinetics of the *paralytic* variants expressed in *Xenopus* oocytes^17^. While there are more protein recoding editing events in flies than in mammals, a number of mammalian ion channels also undergo functionally important RNA editing events, some of which are dynamically regulated across brain tissues in multiple species^18,19^; yet, elucidating the function of a particular editing site may not be fully assessed at the entire tissue level. Editing levels are known to differ between neurons and glial cells ^20^, but little is known about the diversity and the functional importance of this process in different neuronal populations. So far, RNA editing profiling of *Drosophila* neurons faced technical difficulty of reliably defining and isolating certain neuronal populations out of many in sufficient quantity, and thus editing level measurements typically represent an average of editing from large brain regions or whole brain tissue.

Here, we utilized a battery of GAL4 drivers and refined the INTACT method (**I**solation of **N**uclei **T**agged in **A** specific **C**ell **T**ype)^21^ to analyze the spatial distribution of editing events among nine different neuronal populations taken from adult fly brains. To examine the relative editing levels of thousands of known and novel editing sites, we deployed two complementary approaches: RNA-sequencing to quantify editing level measurements in highly expressed transcripts across the different neuronal populations, and microfluidic multiplex PCR (mmPCR) to gain highly accurate editing level measurements at targeted sites^22^. We identified novel editing sites using the RNA-seq data and then determined editing levels at these sites and all previously identified sites though either mmPCR or RNA-seq.

We found that each neuronal population has a unique RNA editing signature composed of distinct editing levels of sites in neuronal proteins, some of which harbor unique combinations of multiple editing sites across the same protein. Many of these regulated sites have been predicted to be functional by previous analyses. We found evidence for co-regulation of nearby sites of the same transcripts and identified instances where different subunits of a certain neuronal machinery are edited differentially in distinct population of neurons. Furthermore, we show that these editing level differences are likely to be caused by factors other than *Adar* expression, suggesting other factors play a role in fine-tuning editing levels across different population of neurons.

## Results

### Isolation of RNA from discrete nuclei-tagged neuronal populations

We and others had previously measured editing levels in whole fly heads and brains^7,23^, but treating the brain as one unit prevented us from pinpointing editing sites that are differentially regulated between distinct populations of neurons, which presumably reflects their functional importance. To reveal RNA editing level variation between types of neurons, we used Gal4 drivers to mark and isolate different subsets of neurons within the fly brain. The chosen neuronal populations regulate various aspects of behavior and physiology and are composed of varying numbers of cells with distinct anatomy and connectivity across the brain (**Figure 1A**). The largest population of cells we studied are the mushroom-body neurons (marked by *OK107-Gal4*)^24^, which serve as the integration center for many behaviors and have a major role in learning and memory^25^, and the Fruitless (*Fru-Gal4*) neurons, which are implicated in specifying social behavior in male and female flies and comprise approximately 2% of central nervous system neurons. We chose four populations of neurons associated with neuropeptide signaling, Neuropeptide-F (*NPF-Gal4*), the Neuropeptide-F Receptor (*NPFR-Gal4*), Diuretic hormone 44 (*Dh44-Gal4*), and Corazonin (*Crz-Gal4*), which regulate different aspects of motivational behaviors and stress response and represent only a small number of neurons in the brain^26^. We also chose three populations expressing neurotransmitters, Dopamine (*TH-Gal4*), Serotonin (*Trh-Gal4*), and Octopamine (*Tdc2-Gal4*), which are implicated in mediating a broad range of innate and learned behaviors as well as regulating homeostatic responses. We also used a pan neuronal driver (*Elav-Gal4*) as a reference for whole brain neurons.

**Figure 1.**
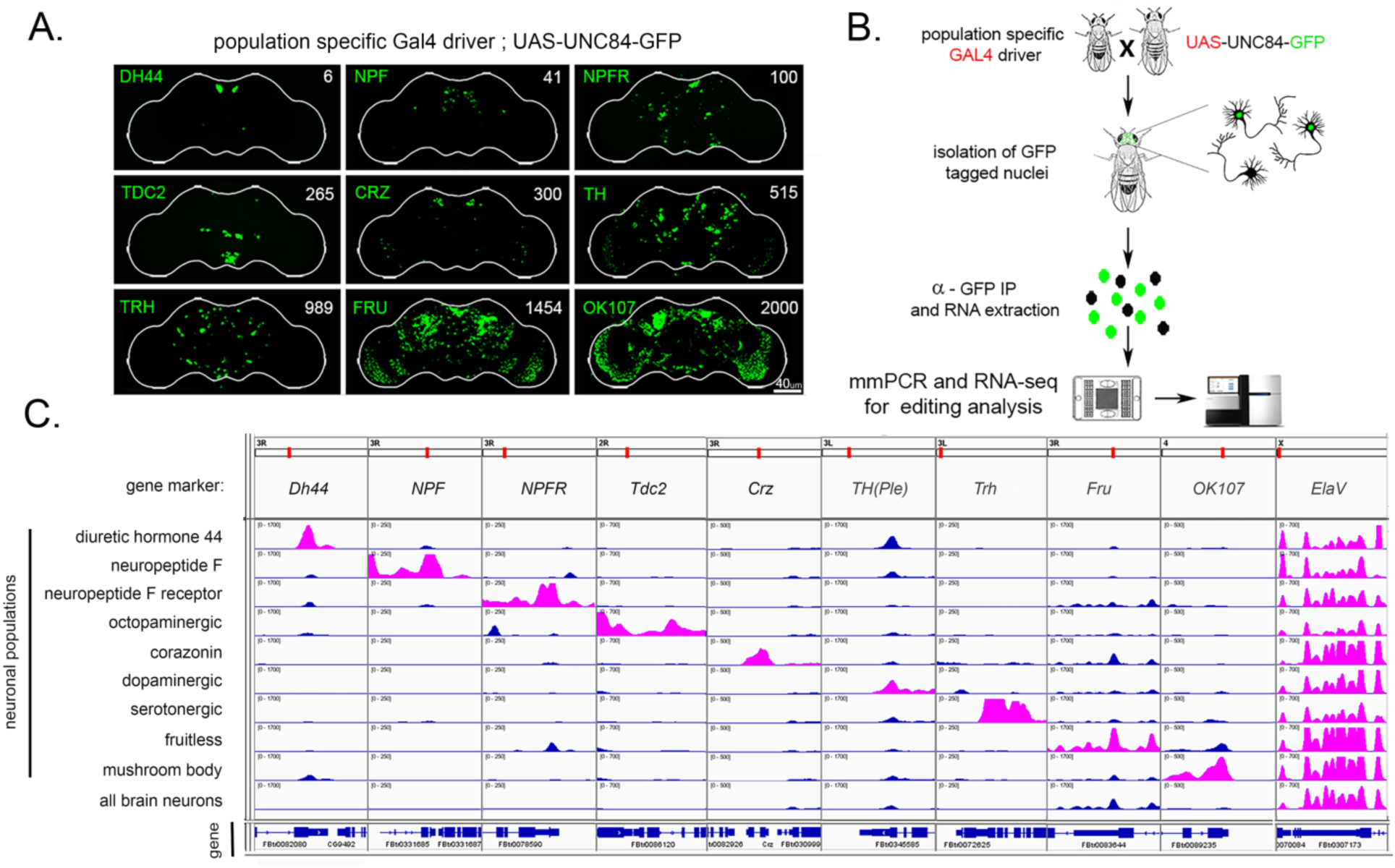
Isolation of RNA from distinct neuronal populations. **(A)** Confocal images of GFP-marked nuclei in fly brains from the nine neuronal populations used in this study. GAL4 drivers are listed on the left, and the number of cells in each neuronal population are listed on the right of each image. Scale bar = 40 um. **(B)** Schematic of workflow for isolating RNA from discrete neuronal populations and RNA editing analysis. **(C)** Visualization of RNA-sequencing reads from the nine populations and Elav control at marker genes for the 10 groups. The reads of the relevant marker genes for each population are listed on left are highlighted in pink.

There are several approaches that allow for the isolation of genetically marked subsets of neurons, including manual sorting^27^, FACS^28^, and ribosome tagging^29^. Since RNA editing is a co-transcriptional process that takes place in the nucleus, we chose to analyze newly formed RNA transcripts residing within neuronal nuclei. For that purpose, we used the INTACT (isolation of nuclei tagged in a specific population) method^21^. This technique utilizes specific *Gal4* drivers to mark neuronal nuclei with a genetically encoded nuclear tag (UNC84-GFP) that can then be purified by immunoprecipitation. Here, we improved upon the previously published INTACT protocol by adding a purification step that minimized nonspecific binding of cytoplasmic debris and fragments of broken nuclei (see **Methods**). We isolated nuclear RNA from 10 specific neuronal populations and used two complementary methods to measure RNA editing across neuronal populations, RNA-sequencing (RNA-seq) and microfluidic multiplex PCR and sequencing (mmPCR-seq)^30^ (**Figure 1B**, see **Methods** for details). We used RNA-seq to measure RNA editing in highly expressed transcripts and mmPCR-seq to obtain highly accurate editing level measurements at 605 loci^31^ harboring known editing sites that did not depend on gene expression.

To validate the capability of our approach in isolating distinct population of neurons, we compared the expression of marker genes across the transcriptome of the 10 different populations of neurons. As expected, most of the marker genes show population specific expression, while the pan neuronal marker *ElaV* is evenly expressed in all groups of neurons (**Figure 1C**). We saw enrichment of the desired markers even for low abundance populations such as NPF or Dh44 neurons, suggesting that we successfully captured the transcriptomes of these neuronal populations. Some neuronal populations, like the one marked by the *TH driver*, had partial overlap with other neuronal populations, as can be seen by the expression of the *TH* marker gene across several neuronal populations.

### Identification of novel sites from distinct neuronal populations

We hypothesized that RNAs from distinct neuronal populations would include novel RNA editing sites that were previously undetected because whole-brain sequencing does not provide adequate coverage of editing sites that are only edited or expressed in a small number of cells. We modified our previously developed computational pipeline to identify novel editing sites from the transcriptomes of the nine neuronal populations as well as the pan neuronal ElaV control (see **Supplementary Note**, **Supplementary Figure 1**). From all populations combined, we identified 2,058 variants of all possible base conversions, 88% of which were A-to-G or T-to-C and thus indicative of A-to-I editing events (**Figure 2A**). These sites included both previously known and novel editing sites in each neuronal population (**Supplementary Table 1**). We identified between 161 (in Crz) and 287 (in Fru) previously known editing sites and 46 (in Dh44) and 518 (in Fru) novel sites in each neuronal population (**Figure 2B**). Many of the novel sites were identified in only one neuronal population, in contrast to the previously known sites, which were more often identified repeatedly in multiple neuronal populations by our pipeline (**Figure 2C**), demonstrating that sequencing each distinct neuronal population facilitated the discovery of additional novel editing sites. The majority of the novel sites we identified did not overlap annotated regions of the transcriptome (**Figure 2D**), and 76% of the novel sites overlapped with annotated repetitive regions of the genome, as compared to 13% of the previously known sites we identified (**Figure 2E**). We found that our novel sites grouped into 225 loci (where there were fewer than 100 bases between adjacent editing sites), with repetitive loci often containing large numbers of sites, including one locus having as many as 116 editing sites (**Figure 2F**).

**Figure 2.**
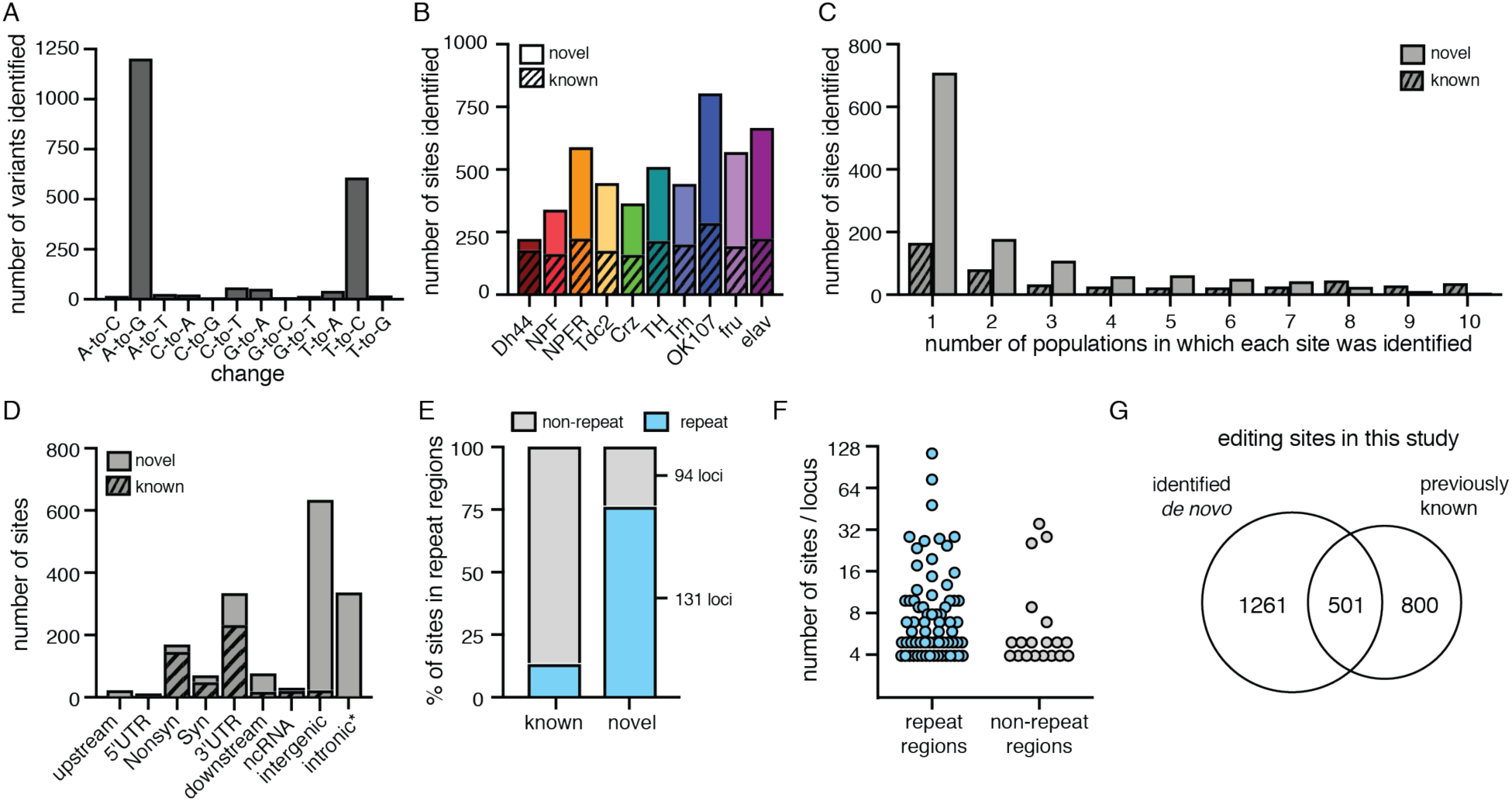
Identification of novel RNA editing sites from distinct neuronal populations. **(A)** The total number of all possible base variants identified *de novo*. **(B)** Stacked bar plot of the number of editing site identified *de novo* from each population with previously known sites in stripes and novel sites in solids. **(C)** Bar plot representing the number of populations in which each known site (striped) and novel site (solid) was identified *de novo*. **(D)** Stacked bar plot of the number of sites found in each annotated location for known (stripes) novel (solid) sites discovered in this work. *The 353 sites annotated as intronic are found in the Myo81F heterochromatic region of chr3R. **(E)** Stacked bar plot representing the percentage of previously known and novel sites identified by our pipeline that overlap annotated repeat regions (blue) or do not (gray). The number of loci containing novel sites is marked for repeat and non-repeat regions. **(F)** Scatter dot plot of the number of editing sites found within each locus that contained at least 4 novel sites for loci overlapping repeat regions and non-repeat regions. Y-axis is log_2_ scale. **(G)** Venn diagram of editing sites identified *de novo* and previously known editing sites used in this study.

In total, we identified 1,762 editing sites using our pipeline (*de novo*), finding 501 previously known editing sites and 1,261 novel sites (**Figure 2G**). Because our *de novo* identification pipeline included stringent filters, for downstream comparative analysis we also measured editing levels at all previously known sites that were highly covered in RNA-seq. We additionally used mmPCR-seq to measure editing levels at previously known sites that were not highly covered in RNA-seq. These two strategies led us to include an additional 800 previously known editing sites in our downstream comparative analysis.

### RNA editing levels differ between neuronal populations in the fly brain

We determined RNA editing levels using both RNA-seq and mmPCR-seq at all previously known editing sites and the novel sites we identified by determining the fraction of G reads over the total number of reads at each site. Both RNA-seq and mmPCR-seq editing level measurements were highly reproducible between three biological replicates from each neuronal population (**Supplementary Figure 2**). In the subset of editing sites that were covered in both the RNA-seq and mmPCR-seq, we saw that editing levels were also highly reproducible between the two methods.

To compare editing levels between neuronal populations, we looked at a total of 1,036 editing sites that were covered by either mmPCR-seq or RNA-seq in at least 7 out of 10 different neuronal populations with 20X coverage and editing levels that were reproducible between replicates (**Supplementary Table 2**). Pairwise comparisons of editing levels between all neuronal populations revealed that 271 editing sites (26% of sites queried) had at least 20% difference in their editing levels between at least two different neuronal populations (**Figure 3A**). To understand which sites were most often different between neuronal populations and which neuronal populations showed the most differences, we counted the number of times the same site was differentially edited between each neuronal population in its pairwise comparisons with all other populations. **Figure 3B** shows the number of sites with decreased or increased editing of at least 20% between each of the ten neuronal populations and every other population as well as the number of the comparisons that show strong differential editing at each site. We found that Fru neurons were the most heavily weighted towards increased editing levels, with 190 editing sites that showed higher editing in Fru than any other neuronal population. Crz neurons were the most skewed towards having lower editing levels than the other neuronal populations, where 155 editing sites had decreased editing compared to at least one other neuronal population (**Figure 3B**). The other neuronal populations showed fewer differentially edited sites than both Fru and Crz, but each of the other populations had between 88 and 160 editing sites that were differentially edited from at least one other neuronal population (**Figure 3B**). We also performed PCA analysis on editing levels in all 10 neuronal populations and observed clustering patterns similar to our pairwise editing comparisons (**Supplementary Figure 3**).

**Figure 3.**
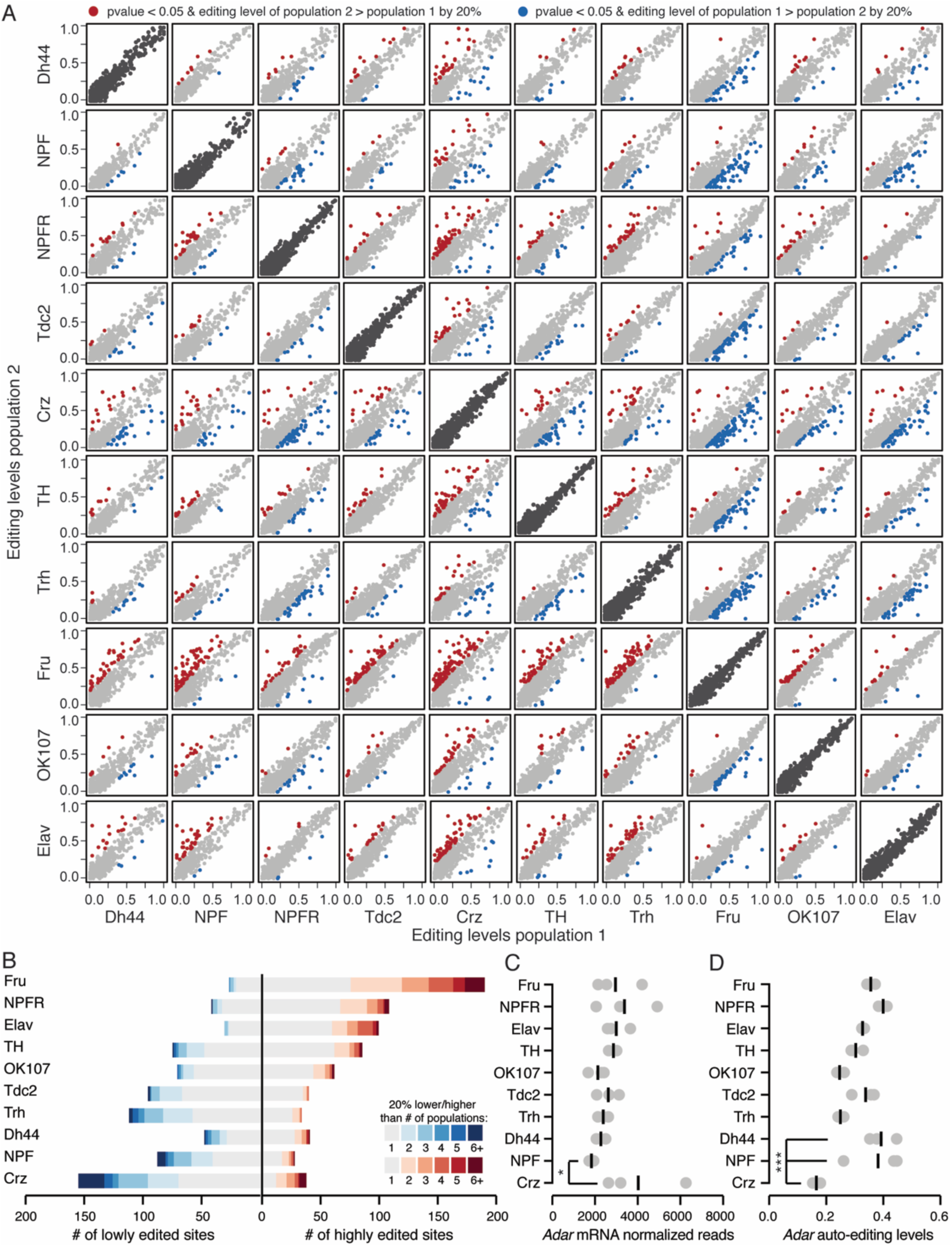
RNA editing level differences between neuronal populations. **(A)** Pairwise comparisons of editing levels from three combined replicates of mmPCR-seq or RNA-seq between 10 populations. Red and blue dots represent editing sites that differ by > 20% editing between populations with p-value < 0.05 by Fisher’s exact test of A and G counts, while gray dots represent sites with < 20% editing between the populations. Dark gray dots are from representative biological replicates of each population. **(B)** The number of editing sites that are more highly or lowly edited in each population listed on the left compared to all other populations. Shades of blue and red represent the number of populations in which each site differs in pairwise comparisons. **(C)** *Adar* mRNA normalized read counts from RNA-seq of each population. Each gray dot is a replicate with black bars representing the mean. * p-value < 0.05 in pairwise comparison by Wald test. **(D)** Editing levels at the *Adar* auto-editing site at chrX:1781840 in all populations. Each gray dot is one replicate, with black bar representing the mean. *** difference in editing > 20% and p-value < 0.001 from Fisher’s exact test.

To test whether the differences in editing levels between these neuronal populations stem from variation in ADAR levels, we determined *Adar* mRNA expression levels in these populations (**Figure 3C**). We found that *Adar* levels were similar between neuronal populations, and that there was no correlation between *Adar* expression and the observed differences in overall editing levels between the different neuronal populations (**Figure 3C and Supplementary Figure 4A**). Next, we tested whether the differences between the populations could be explained by variation in the ADAR enzymatic activity, which is decreased upon auto-editing within its own transcript (at chrX:1781840)^13^. Like *Adar* expression, auto-editing levels were similar between neuronal populations, with Crz neurons having the lowest auto-editing levels of all neuronal populations (**Figure 3D and Supplementary Figure 4B**). This lower auto-editing level would be expected to contribute to increased, rather than decreased editing levels in Crz; therefore, we concluded that auto-editing levels are not mainly responsible for editing differences between neuronal populations. Furthermore, we found that there was no correlation between the number of cells in each neuronal population and overall editing levels (**Supplementary Figure 4C**).

### Identifying unique regulation of editing events in different neuronal populations

We then sought to identify population-specific “outlier” sites that were differentially regulated in one neuronal population compared to all others. We calculated z-scores to determine how much each replicate of each population of neurons differed from the mean of all population replicates at each site (**Supplementary Table 3**). We identified 31 editing sites that were lowly edited in one population (**Figure 4A**) and 33 sites that were highly edited in one population (**Figure 4B**). The majority of both the lowly and the highly edited sites are found in Crz and Fru neurons, respectively, consistent with our previous analysis (see Figure 3B). To further characterize the list of population-specific editing differences, we performed gene ontology (GO) analysis of the genes containing these differentially edited sites. We found that Crz-specific editing differences are located within genes enriched for multiple GO terms related to the regulation of membrane potential and cation transmembrane transport above a background of edited genes in this dataset (**Supplemental Table 4**). Fru-specific editing sites were found more often in transcripts that play a role in cell signaling and differentiation, but they did not show any statistically significant GO term enrichment.

**Figure 4.**
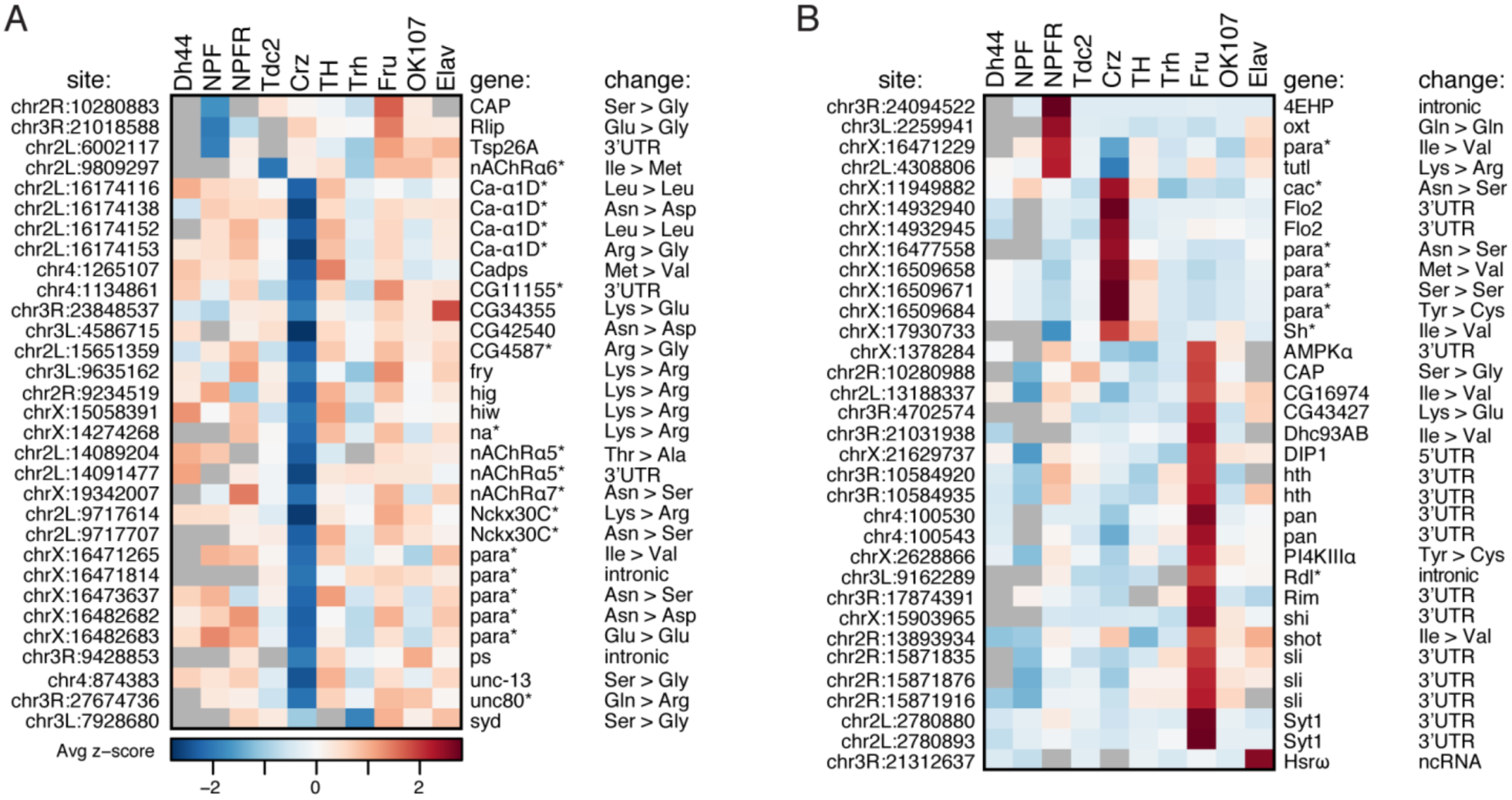
Population-specific editing level differences. **(A-B)** Average z-score of replicate editing levels at sites where one population shows a population-specific decrease in editing **(A)** or a population-specific increase in editing **(B)**. Genes with (*) are involved in ion transport.

While editing differences between tissues have previously been associated with expression differences of edited transcripts^32^, the majority of transcripts with population-specific editing did not show differential expression in the neuronal population in which they were uniquely edited (**Figure S5**). However, five transcripts that are edited differently in Crz neurons are also expressed more highly in Crz neurons than all of the other neuronal populations: *CG34355*, *Flo2*, *para*, *nAChRalpha7*, and *Nckx30C*. Nevertheless, these transcripts that are highly expressed in Crz contain instances of both decreased and increased editing levels, implying that the site-specific differential editing levels cannot be simply explained by the relative abundance of the transcript within a specific neuronal population.

### Co-regulation of clustered editing sites

Of the 64 sites that had population-specific editing levels, we found a total of 19 sites in 8 different groups that were located within about 40 bases of at least one other population-specific site. Since nearby editing events in *Drosophila* often occur within the same physical transcript^33^, we examined whether these groups of sites showed similar population-specific editing trends due to co-regulation of the editing events in the clusters. We measured the usage of each possible editing isoform in all neuronal populations for each cluster of sites that could be found within the same amplicon from mmPCR-seq. First, we looked at a cluster of two sites in the *Sli* transcript that are 40 bases apart and both highly edited in Fru neurons. These sites are edited at 29% and 34% in Fru, whereas the median editing levels across all populations are 10% and 18% respectively (**Figure 5A**). We calculated the usage of the four possible editing isoforms covering these two sites, and we found that in Fru, the AA isoform was used less often than in the other populations (67% compared to the median 84%) and the GG isoform was used more often in than the other populations (26% compared to the median 10%) (**Figure 5B**). Based on the editing levels measured at the two sites independently, we would expect the AA and GG isoforms to represent 50% and 8% of the total number of transcripts, but we found both the AA and GG to be over-represented by 17%, with a concomitant decrease in AG and GA transcripts (**Figure 5C**), suggesting that the editing events at the two sites are in fact linked. We then measured the differences in the observed versus the expected percentages of isoform usage in the 5 other clusters of two sites that appeared to be co-regulated for all populations. We found that, in all populations, these sites also showed an over-representation of AA and GG isoforms and an under-representation of AG and GA isoforms (**Figure 5D** and **Supplementary Figure 6A-E**), confirming that in these clusters ADAR preferentially edits either both sites together or neither site.

**Figure 5.**
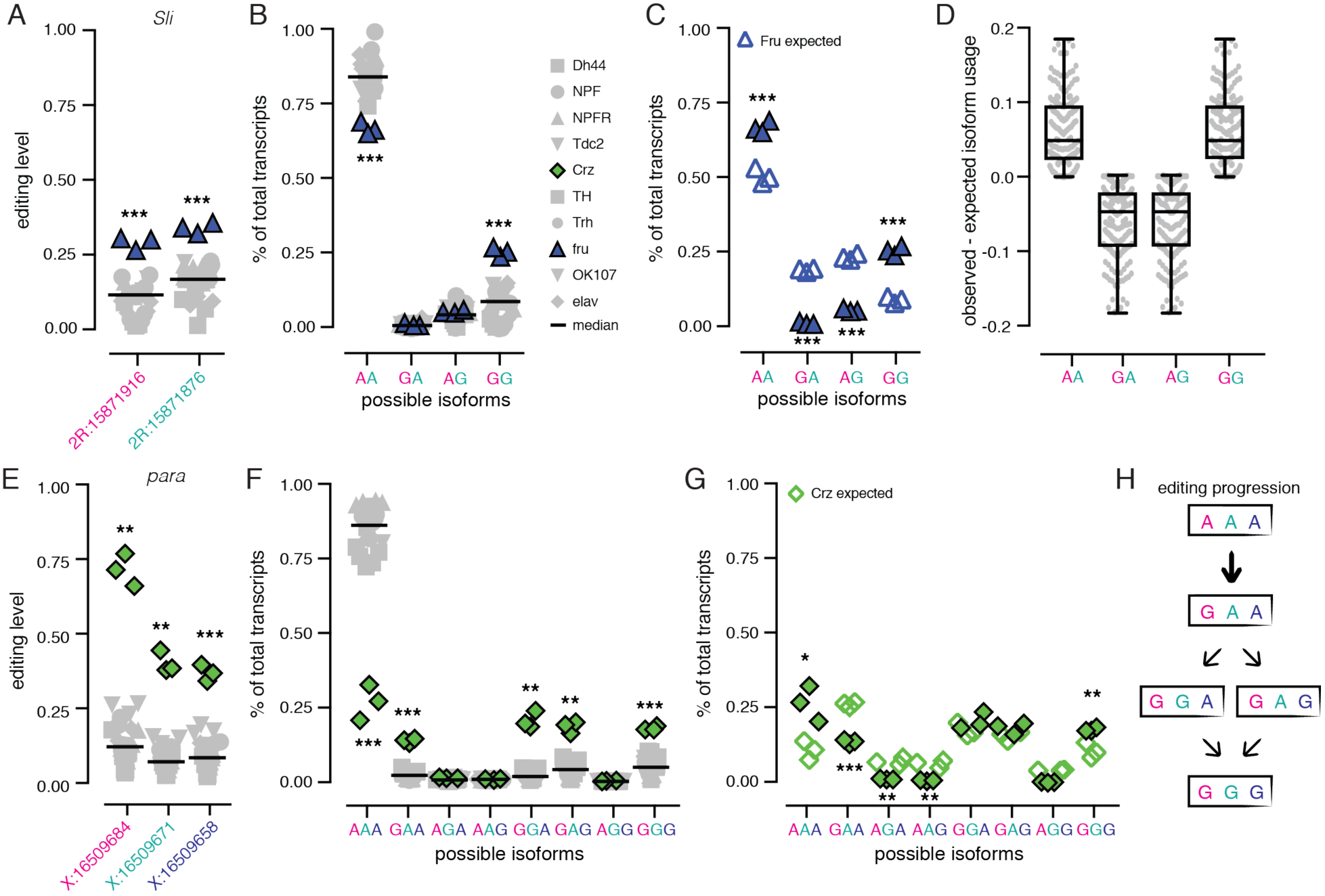
Co-regulation of proximal editing sites. **(A)** Editing levels across all populations at a cluster of two editing sites in *Sli*. Fru is highlighted in blue. *** p-value < 0.001 by Welch’s t-tests. **(B)** The percentage of total transcripts using each possible editing isoform at the two sites in all populations. Fru is highlighted in blue. *** p-value < 0.001 by Welch’s t-tests and mean difference > 10%. **(C)** Observed and expected isoform usage in Fru neurons. *** p-value < 0.001 by Student’s t-tests. **(D)** Boxplots depicting the difference between the observed and expected isoform usage of four isoforms for six clusters of co-regulated sites in all populations. **(E)** Editing levels across all populations at a cluster of three editing sites in *para*. Crz is highlighted in green. ** p-value < 0.01, *** p-value < 0.001 by Welch’s t-test. **(F)** The percentage of total transcripts using each possible editing isoform at the three sites in all populations. Crz is highlighted in green. ** p-value < 0.01, *** p-value < 0.001 by Welch’s t-tests with mean difference > 10%. **(G)** The observed and expected isoform usage in Crz neurons. * p-value < 0.05, ** p-value < 0.01, *** p-value < 0.001 by Students t-tests. **(H)** A model for editing at the cluster of three sites shows that editing at the first site is critical for editing at the other sites in the cluster.

In addition to the co-regulated clusters of two sites, we also identified two larger clusters of sites that showed similar evidence of co-regulation, including a cluster of four sites in ca-alpha1D (**Supplementary Figure 6F**) and a cluster of three sites in *para*. This three-site cluster showed editing increases of 58%, 32% and 27% over the median editing levels of the other neuronal populations (**Figure 5E**). We found that, at the isoform level, these editing increases at the three sites lead to a 59% decrease in completely unedited transcripts (AAA) from the median level of all populations and a similar increase (between 12% and 19%) of four different editing isoforms: GAA, GGA, GAG, and GGG (**Fig 5F**). Similar to the clusters containing two sites, we found that the completely unedited isoform and the completely edited isoform (AAA and GGG) were over-represented, while isoforms with only one editing event (GAA, AGA, AAG) were under-represented (**Figure 5G**). From this data, we can postulate a progression of editing at these three sites. Since all edited isoforms included editing at the first site, we propose that this site is edited first, and that editing at one or both of the second and third sites in the cluster requires editing of the first site (**Figure 5H**).

### Differential RNA editing in transcripts involved in neuronal transmission

A substantial proportion of the transcripts with population-specific editing are known to play a role in neuronal transmission, so we wanted to explore the consequences of these editing differences. We compared our population-specific editing sites to three recent studies that used computational strategies to predict editing events in *Drosophila* that are likely to be functional because they are found in conserved regions of the genome, are conserved as editing events throughout multiple *Drosophila* species, or are in regions under positive selection^9-11^. Of the 64 population-specific editing sites, 41 sites were predicted to be likely functional by at least one of these studies, while only 8 sites showed evidence against functionality (the remaining 15 were not studied, see **Supplementary Table 2**).

Of the sites that are predicted to be functional, many are found in transcripts that encode proteins that are critical for neuronal function, and some are known to function together within the same multiprotein complex or in the same pathway. In Crz neurons, Ca-alpha1D, which encodes an alpha subunit of a voltage-gated calcium channel, had two editing sites that showed a decrease in editing of 26% and 42% from median editing levels across all neuronal populations (Figure 6A). While these editing decreases may alter the function of this calcium-gated ion channel, they are not the only editing differences in Crz neurons that might have an effect on calcium ion flow. We also observed an editing increase of 27% at a site within the EF-hand calcium-sensing domain of the voltage-gated calcium channel, cacophony (cac). Furthermore, we observed a decrease in editing of 22% from the median level of all populations in CG4587, which encodes an auxiliary α2δ subunit of voltage-gated calcium channels (**Figure 6A**). The population-specific editing of the different subunits suggests that each neuronal population possesses a unique mixture of voltage-gated calcium channel isoforms that is the result of the distribution of the different subunit combinations. We also observed a similar regulation of editing at putatively functional sites in two transcripts encoding the sodium leak channel complex. One editing site within each of the narrow abdomen (na) α subunit and its auxiliary subunit Unc80^34^ showed decreased editing of 19% in Crz neurons from the median levels of all populations (**Figure 6B**). Various acetylcholine-gated ion channels (nicotinic acetylcholine receptors; nAChRs) also showed editing differences between populations. nAChRalpha6 and nAChRalpha7 showed decreases in editing in Tdc2 of 27% and in Crz of 21%, respectively, at sites that change amino acids in the ligand-binding domains of these related proteins, while nAChRalpha5 showed 40% lower editing in Crz at one site predicted to change an amino acid in the ion-channel pore domain (**Figure 6C**). The identification of differential editing levels of several protein subunits that function together highlights the strength of RNA editing in diversifying the proteomic architecture. The composition of neuronal machineries within a certain neuronal population is determined by the distribution of edited versus non-edited forms of a given subunit and its association with other subunits that undergo differential editing and function within the same multiprotein complex. These data may indicate co-regulation of editing across related transcripts in different neuronal populations.

**Figure 6.**
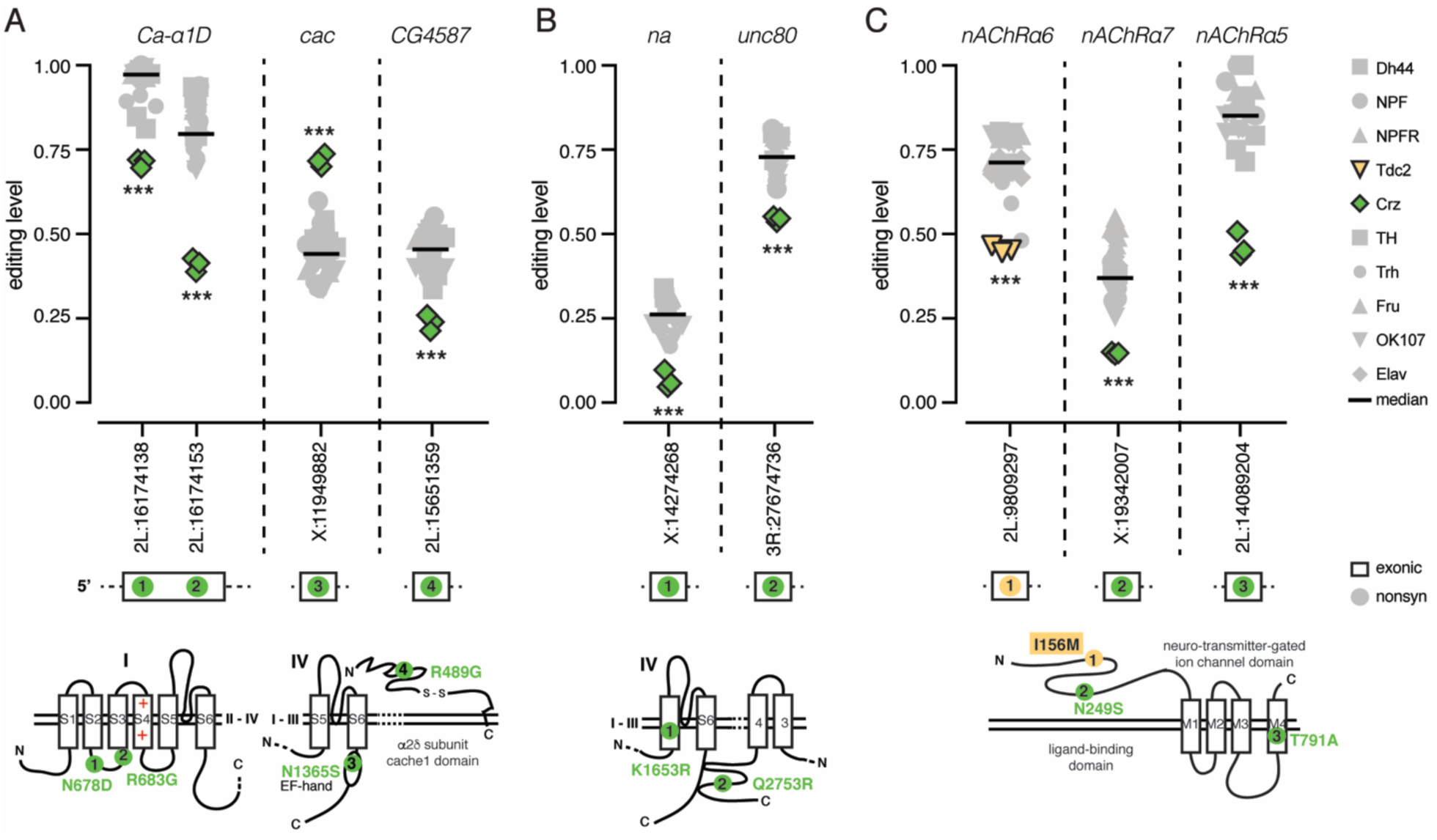
Co-regulation of RNA editing in related proteins. **(A)** RNA editing levels at sites that alter amino acid sequences within calcium-gated ion channel subunits, Ca-alpha1D, cac, and CG4587. Crz editing is in green. **(B)** RNA editing levels at sites that alter amino acid sequences within sodium leak channel components, na and Unc80. Crz editing is in green. **(C)** RNA editing levels at sites that alter amino acid sequences within nicotinic acetylcholine receptor subunits, nAChrRalpha5, nAChrRalpha6, and nAChrRalpha7. Tdc2 editing in nAChRalpha6 is in yellow, Crz editing in nAChRalpha5 and nAChRalpha7 is in green, with other populations in gray. Black bars represent median editing of all populations. * p-value < 0.05, ** p-value < 0.01, *** p-value < 0.001 from Welch’s t-test. Site locations are noted as chromosome: position. Cartoon protein structures shows amino acid location of sites that cause nonsynonymous changes, numbered in cartoon transcripts above. See Supplementary Table 7 for protein annotation information.

### Editing differences suggest functional differences

Numerous editing sites that showed differential editing levels between neuronal populations occurred within voltage-gated ion channels Para and Shaker. Voltage gated ion channels are composed of four subunits or four linked subunit-like repeats, each of which contains six transmembrane segments (S1-S6)^35^. In Crz neurons, the transcript encoding the voltage-gated sodium channel Para was differentially edited at 10 different sites along its transcript, with 6 sites showing decreased editing and 4 sites showing increased editing levels. The first site, changing Tyr^189^ to Cys shows a remarkable 58% editing increase in Crz neurons over the median level in other neuronal populations, while the last site, an Ile^1691^ to Val change in the S3 of repeat IV showed a 45% increase in NPFR neurons (**Figure 7A**). The Tyr^189^ in the S2 of the voltage-sensing domain (VSD) of repeat I is highly conserved across different species (**Figure 7B**), leading us to hypothesize that an editing change might alter protein function.

**Figure 7.**
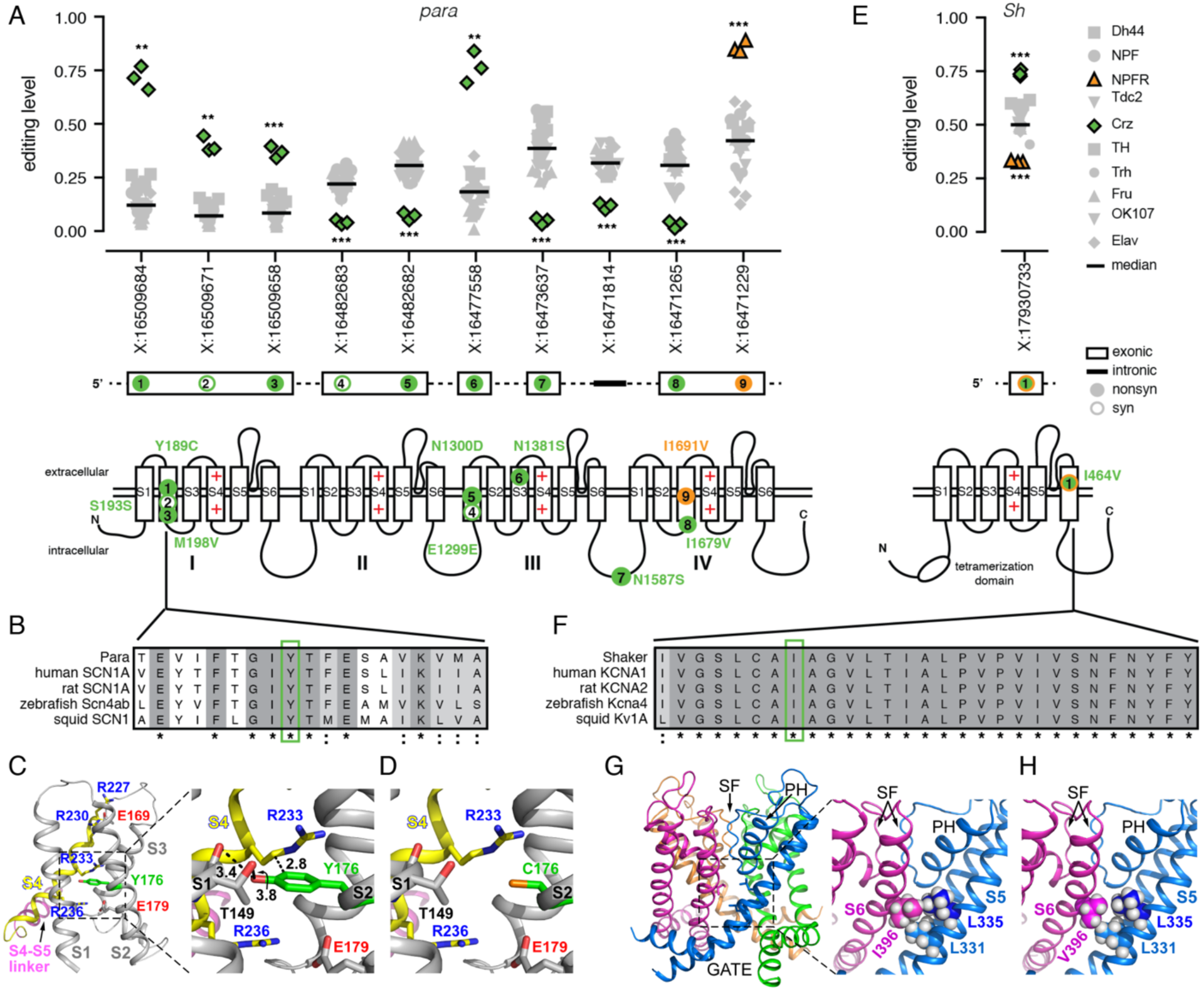
RNA editing in voltage-gated ion channels Para and Sh. **(A)** RNA editing levels at population-specific editing sites in *paralytic*. Crz is in green, NPFR is in orange for significant sites and all other populations are gray. Black bar represents median editing levels of all populations. * p-value < 0.05, ** p-value < 0.01, *** p-value < 0.001 from Welch’s t-test. Location of amino acids changed by nonsynonymous sites (filled circle) and synonymous sites (open circle) in the protein are marked on cartoon protein structures. **(B)** Amino acid conservation within the S2 transmembrane domain of repeat I across voltage-gated sodium channels of five species with Tyr^189^ highlighted in green. **(C)** The Tyr189Cys amino acid change in *Para* mapped onto the 3-D structure of the homologous voltage-gated Na^+^ channel from *Periplaneta Americana*. Ribbon diagram of the voltage sensor domain (side view of S1-S4). Tyr^176^ (green) is homologous to *D. melanogaster* Tyr^189^. Magnification (right) showing the potential interactions (dashed lines) of Tyr^176^ (in S2) with Thr^149^ (in S1) and Arg^233^ (in S4). **(D)** Same as in C, but reflecting the RNA editing of Tyr^176^ to Cys. Carbon atoms are shown as gray or yellow ribbons. Oxygen, nitrogen, and sulfur atoms are shown in red, blue, and orange, respectively. Numbers near dashed lines indicate distances in angstroms. Hydrogen atoms were removed for clarity. **(E)** RNA editing levels at an editing site with population-specific editing in *Shaker*. **(F)** Amino acid conservation within the S6 transmembrane domain across voltage-gated potassium channels of five species with Ile^464^ highlighted in green. **(G)** The edited amino acid Ile464Val of Shaker mapped onto the 3-D structure of the *Rattus norvegicus* Kv1.2 voltage-gated K^+^ channel. Ribbon diagram of the pore domain of Kv1.2 as seen from the side, with the four identical subunits colored differently. SF, selectivity filter. PH, pore helix. Magnification (right) showing Ile^396^ (homologous to Ile^464^) of the S6 segment and its potential van-der-Waals interactions with Leu^331^ and Leu^335^ of S5 in the adjacent subunit. **(H)** Same as in panel G, but reflecting the RNA editing of Ile^396^ to Val.

To predict the functional consequences of recoding Tyr^189^ into a Cysteine residue (Y189C), we mapped the edited amino acid onto the 3-D structure of the homologous voltage-gated sodium channel from American cockroach (PDB ID 5X0M) (**Figure 7C-D**). Like in other voltage-gated ion channels, the S2 segment is part of the VSD formed by the first four transmembrane segments, S1-S4; whereas the fifth and sixth transmembrane segments (S5 and S6) are tightly arranged around a four-fold axis of symmetry to create the ion conduction pathway^36-39^ (**Figure 7A** lower panel). VSDs of various voltage-dependent ion channels are endowed with charged amino acids, also called gating charges, and have four highly conserved arginine residues along S4 (R1, R2, R3, and R4)(e.g., **Figure 7C**) that mainly contribute to the voltage-driven gating charge transfer during channel activation^40-46^. The gating charges reside in aqueous crevices and translocate across a focused electric field that is occluded by a bulky residue (Phe or Tyr)^37-39^. It has been suggested that the charge transfer across the Phe/Tyr bulky residue on S2 is facilitated through sequential electrostatic interactions of the gating charge residues with negative countercharges in segments S2 and S3^47-50^. Inspection of the 3-D structure indicates that recoding Tyr^189^ into Cys (homologous to the aforementioned Tyr^176^ in PDB ID 5X0M) would eliminate the bulky occlusion in the pathway of the gating charges in the first VSD (**Figure 7C-D**), which might modify gating kinetics.

Crz and NPFR neurons also showed differential editing at a site in the voltage-gated potassium channel Shaker (Sh) that is predicted to change Ile^464^ to Val (I464V) in the S6 pore-lining segment. The site shows a 24% increase in editing in Crz neurons and 17% decrease in NPFR neurons compared to the median level of all populations (**Figure 7E**). Ile^464^ and its flanking amino acids are highly conserved among potassium channels (**Figure 7F**). To assess the structural effect of such an RNA editing event, we inspected the X-ray crystal structure of the Kv1.2 voltage-dependent K^+^ channel (PDB ID 2A79)^51^, a homologous potassium channel from *Rattus norvegicus*. The 3-D structure of the rat Kv1.2 indicates that its Ile^396^ of the S6 pore-lining segment (homologous to Ile^464^ in the *Drosophila* shaker K^+^ channel) forms van der Waals interactions with Leu^331^ and Leu^335^ on the S5 segment of the adjacent subunit (**Fig 7G-H**). One can therefore envision that the replacement of Ile^396^ by valine in the Kv1.2 structure might result in the loss of the van der Waals interaction with Leu^335^ (**Fig 7G-H**) and weaken the bond network between S6 and S5. Such a structural perturbation might propagate to the pore helix and the selectivity filter and therefore affect C-type inactivation and/or might propagate along the S6 towards the activation/deactivation gate.

## Discussion

We set out to increase the resolution of our understanding of the transcriptome-wide RNA editing landscape within the brain by determining editing level differences between nine different neuronal populations in the fly. By improving the INTACT protocol, we were able to isolate different populations of neurons with distinct functional differences. Using RNA-seq and mmPCR-seq, we determined how RNA editing plays a role in facilitating transcriptomic diversity between these functionally diverse populations.

We identified many novel editing sites in these neuronal populations. These novel sites, which were often found in lowly expressed transcripts, mostly overlapped repetitive regions of the transcriptome and contained numerous editing sites, which is consistent with previous reports of editing of repeat regions in flies and many other species including Alu sequences in human^52,53^. While most studies of RNA editing in *Drosophila* have focused primarily on ADAR editing of coding regions, our data suggest that ADAR has a wide-ranging role in editing non-coding transcripts. These editing events may regulate transposable elements, circRNA biogenesis54, and RNA interference pathways, which can in turn alter heterochromatin formation^52^. They may also play a role in the *Drosophila* innate immune system, distinguishing self from non-self RNAs, similar to demonstrated roles for ADAR proteins in mammals^16,63^. While additional studies are needed to determine the functional significance of these editing events, our data suggest that sequencing the transcriptomes of small neuronal populations can facilitate the discovery of these sites by providing deep sequencing of rare RNAs.

We identified hundreds of sites where editing differed between at least two groups of profiled neurons. In contrast to a previous report that showed high editing levels mainly in mushroom body neurons using a reporter construct of an engineered editing substrate^55^, we found prominent editing across all of these populations of neurons, with editing levels similar to mushroom body neurons. The editing differences we identified were found at specific sites, rather than being global changes to editing at all sites. Therefore, these differences could not be explained by ADAR expression or auto-editing differences between populations. The transcripts containing the differentially edited sites were also generally similarly expressed, suggesting a complex regulation of editing levels at these sites.

Our results indicate that Crz and Fru neurons stand out as particularly different from the other neuronal populations studied in terms of their editing signatures. In some transcripts, we found bi-directional regulation of editing across the same transcript. We also identified a number of editing sites that were physically close and co-regulated in the same neuronal populations, as ADAR is likely to edit these groups of sites sequentially. These types of editing differences suggest a regulation of editing that can exert its effect differently in different parts of the same transcript. Regulation by RNA binding proteins may have such an effect^56^, but the RNA binding proteins known to regulate editing in flies, the Fragile X protein^57^ and the RNA helicase Maleless^58^ cannot explain the editing differences we observed, suggesting other regulators are yet to be discovered.

In addition to RNA binding proteins, the circadian clock gene *period* has been shown to influence RNA editing in flies^59^. Flies with hypomorphic alleles of *Adar* also show defects in circadian rhythm^60^, suggesting a connection between RNA editing and circadian rhythm. Interestingly, some Crz neurons have been shown to express *period*, signifying Crz neurons may play a role in circadian rhythm^61^, which could be important for regulation of the editing differences that we observe in Crz neurons. In fact, two related proteins that we found to be differentially edited in Crz neurons, Unc80 and na, are known to play critical roles in circadian rhythm^34^. Further functional study is needed to fully determine whether RNA editing in Crz neurons in particular contributes to circadian rhythm.

A number of the differentially regulated sites we identified across these neuronal populations were predicted to be functional in recent computational analyses of RNA editing in *Drosophila*^9-11^, suggesting that the editing differences we observe between neurons may have physiological consequences for the fly. Based on high homologies with ion channels having resolved 3-D structures, we predict that editing of two sites may alter voltage sensing and gating kinetics in the Paralytic sodium channel and the Shaker potassium channel, presumably leading to functional differences in neuronal excitability or sensitivity to different neuromodulators. The Ile^464^ to Val editing in the *Drosophila Shaker* K^+^ channel is lowly edited in NPFR neurons and highly edited in Crz neurons. Previous electrophysiological studies showed that, when N-type inactivation is removed, the V464-edited isoform of the *Drosophila* Shaker K^+^ channel displays a significantly slower deactivation rate than the I464-unedited channel^14^. A more physiologically interesting effect emerged when N-type inactivation was characterized in the wild type edited and unedited isoforms of the Shaker K+ channel^14^. That is, compared to the I464-unedited isoform, the V464-edited isoform inactivates more rapidly (significant between -25 and -15 mV) and displays stronger steady-state inactivation and recovers more slowly from inactivation^14^. Such alterations in N-type inactivation would likely lead to broadening of the action potentials, as is the case when voltage-gated K^+^ channels in rat mossy fibers inactivate rapidly and recover from inactivation very slowly^62^. In fact, editing events nearby, such as the editing site that changes Ile^470^ to Val and is conserved in humans, have profound effects on protein function^19,63^. We therefore hypothesize that the excitability of Crz neurons in the *Drosophila* changes upon I464 to V editing in the Shaker K^+^ channel. Crz neurons also show increased editing at Tyr^189^ in the S2 segment of repeat I of the Para channel. Based on 3-D modeling, we predict that this change might alter gating kinetics by altering amino acid interactions within the voltage-sensing domain of the protein; however, whether editing would confer different gating properties on the channel remains to be elucidated experimentally. In addition to protein-recoding differences, we also observe editing changes in 3’UTRs, which our previous analysis predicts can be functional^9^ and might alter gene expression, mRNA localization, or other post-transcriptional regulatory mechanisms. The functional insights provided by our study can prompt future in-depth biochemical and behavioral analysis that were previously hampered by the need to choose which of the thousands of editing sites to focus on. The identification of highly regulated sites and their spatial distribution across different neurons can promote studies to dissect their functional relevance within the right cellular context.

One caveat in considering the functional consequences of these editing sites is taking into account other sites within the same protein that might also alter protein function. We measured editing isoforms for a set of sites that appeared to be co-regulated in different neuronal populations, and we found that editing sites that reside within 40 bases of each other in a transcript were often edited in tandem in the same physical transcript. We show here that the Tyr^189^ event in para is closely linked with another non-synonymous amino acid change as well as a synonymous change. Since editing events in Shaker have been shown to display functional epistasis^14^, this linkage of editing may enhance or attenuate functional consequences of the editing event. We also show that editing of related proteins, such as subunits of voltage-gated calcium channels, can show co-regulation within neuronal populations, which might create greater functional differences between these populations.

Decreasing editing at many sites by knocking down ADAR in a number of different neuronal populations leads to locomotor and behavioral changes in the fly^60,64^; however, it is unclear whether the regulation of specific editing sites contribute to these behavioral changes, or that it reflects global impairment of neuronal function due to dysregulation of many targets. The data presented in this study, serve as a valuable resource towards identifying functionally relevant editing events, as it exposes highly regulated sites which can serve to bridge the gap between their cell-specific function and regulation of complex behaviors.

## Methods

### Fly stocks and culture

Flies were raised at 25°C in a 12-h light/12-h dark cycle in 60% relative humidity and maintained on cornmeal, yeast, molasses, and agar medium. *UAS_-_unc84-2XGFP* transgenic flies were crossed with the following GAL4 driver: *Dh44-GAL4, NPF-GAL4, NPFR-GAL4, Tdc2-GAL4, CRZ-GAL4, TH-GAL4, TRH-GAL4, Fru-GAL4, OK107-GAL4 and ElaV-GAL4. NPFR-GAL4* was a gift from the Truman lab (HHMI Janelia Campus).

### RNA extractions from different neuronal populations

Neuronal population specific labeled nuclei were isolated using the INTACT method (Isolation of Nuclei Tagged in A specific Cell Type technique) as previously described^1^. This method was slightly modified as follows: about 300 adult flies collected from 2-3 days F1 generation of 10 different GAL4 driver X *UAS_unc84_2XGFP*_reporter was anesthetized by CO^2^ and flash frozen in liquid N_2_. Heads were separated by vigorous vertexing followed by separation over dry-ice cooled sieves. 9ml of homogenization buffer (20mM β-Glycerophosphate pH7, 200mM NaCl, 2mM EDTA, 0.5% NP40 supplemented with RNAase inhibitor,10mg/ml t-RNA, 50mg/ml ultrapure BSA, 0.5mM Spermidine, 0.15mM Spermine and 140ul of carboxyl Dynabeads -270 Invitrogene: 14305D) was added to each sample. The heads were minced on ice by a series of mechanical grinding steps followed by filtering the homogenate using a 10um Partek filter assembly (Partek: 0400422314). After removing the carboxyl-coated Dynabeads using a magnet, the homogenate was filtered using a 1um pluriSelect filter (pluriSelect: 435000103). The liquid phase was carefully placed on a 40% optiprep cushion layer and centrifuged in a 4°C centrifuge for 30min at ~2300Xg. The homogenate/Optiprep interface was incubated with anti-GFP antibody (Invitrogen: G10362) and protein G Dynabeads (Invitrogen: 100-03D) for 40 minutes at 4°C. Beads were then washed once in NUN buffer (20mM β-Glycerophosphate pH7, 300mM NaCl, 1M Urea, 0.5% NP40, 2mM EDTA, 0.5mM Spermidine, 0.15mM Spermine, 1mM DTT, 1X Complete protease inhibitor, 0.075mg/ml Yeast torula RNA, 0.05Units/ul Superasin). Bead-bound nuclei were separated using a magnet stand and resuspended in 100ul of RNA extraction buffer (Picopure kit, Invitrogen # KIT0204), and RNA was extracted using the standard protocol.

### mmPCR-seq

We performed mmPCR-seq^22^ to quantify editing levels at 605 loci harboring know editing sites. We prepared samples for microfluidic PCR with a 15-cycle pre-amplification PCR reaction using 10 ul of cDNA made from INTACT RNA extractions, using the High capacity cDNA Reverse Transcriptase Kit, 6 ul of a pool of all primers used in the multiplex microfluidic PCR, and 4 ul of 5X KAPA2G Fast Multiplex. The pre-amplification reactions were purified using AMPure XP PCR purification beads (Beckman Coulter). We loaded the pre-amplified samples and 48 pools of PCR primers designed to amplify Drosophila editing sites of interest ^31^, into a 48.48 Access Array IFC (Fluidigm) and performed target amplication as previously described^30^. Multiplex PCR products were barcoded using a 13-cycle PCR reaction. After barcoding reaction, samples were pooled and purified using AMPure XP PCR purification beads and were sequenced using Illumina NextSeq with paired-end 76 base pair reads.

### RNA-seq library preparation and sequencing

The NuGEN RNAseq v2 (7102-32) kit was used to prepare cDNA from the INTACT purified RNA, followed by library preparation using the SPIA - NuGEN Encore Rapid DR prep kit. Samples were sequenced on an Illumina HiSeq using single-end 60 base pair reads.

### Identification of novel editing sites

To identify novel sites from each neuronal population, we merged RNA-sequencing reads from three replicates of each neuronal population together as input to our pipeline. We also merged all replicates of all neuronal populations as an additional input to identify novel sites. RNA-sequencing reads were mapped to the dm6 (Aug 2014, BDGP Release 6 + ISO1 MT/dm6)^65^ genome using STAR (v2.4.2)^66^ (--twopassMode Basic) after trimming low quality bases using Trim Galore. Mapped reads were processed using GATK (v3.6)^67^ for indel realignment and duplicate removal and to call variants. We removed variants that overlapped known SNPs from the DGRP^68^, dbSNP, and a recent study^69^, and variants found at the beginning of reads, near splice junctions, in simple repeat regions, or in homopolymeric runs as described in ^7^. We further filtered variants to remove those with less than 10X coverage, less than 10% editing level, or fewer than 3 alternative nucleotides. We then required variants to be present in at least two of three biological replicates. Finally, because we know *bona fide* editing sites are often found in clusters, we removed any variants that were not found next to a variant of the same type in the same transcript. For example, if the nearest variants to an A-to-G change were C-to-T and G-to-A, we would discard the A-to-G, but if one of the adjacent variants was instead an A-to-G it would be kept. This pipeline produced novel editing sites with a false discovery rate of 9%.

### Determining editing levels from mmPCR and RNA-seq

To determine editing levels at known and novel sites, we used STAR (v2.4.2)^66^ (--twopassMode Basic) to map paired-end mmPCR-seq reads and single-end RNA-seq reads to the dm6 genome as described above. We then used the Samtools ^70^ mpileup function to determine base calls from uniquely mapped reads at known and novel editing sites, and calculated editing levels as number of G reads divided by the total of both A and G reads at a site. For mmPCR-seq, we required each replicate to have 100X coverage and we removed sites that were not within 20% editing between replicates, as done previously^31^. Final mmPCR editing levels were determined after down sampling coverage to 200 reads for statistical analysis. For RNA-seq, we required 20X coverage from non-duplicate reads. The majority of sites had either mmPCR-seq or RNA-seq coverage. If we had both mmPCR and RNA-seq coverage at the same editing site, we used the data from mmPCR-seq only. Differences between editing levels were then determined using Fisher’s exacts comparing A and G counts from one sample to another, with a multiple hypothesis testing correction with p.adjust() using a Benjamini and Hochberg correction^71^ Corrected p-values < 0.05 were considered significant. Statistical tests were performed using R (v3.4.1).

### Determining population-specific editing events

We called editing sites population-specific if the absolute values of the z-scores for all replicates for one neuronal population were greater than 1.65 and the editing level of that neuronal population was at least 10% different from the next closest population

### Editing site annotations

Editing sites were annotated using RefSeq gene annotations and ANNOVAR software^72^. To determine protein changes as a result of these editing changes, we used protein annotations from Uniprot^73^. RefSeq and Uniprot ID numbers can for highlighted transcripts can be found in **Supplementary Table 7**.

### GO Term analysis

We used Flymine^74^ to determine GO term enrichment from a background gene set that included all genes with editing sites that were used in comparative editing analysis.

### Determining gene expression levels from RNA-seq

Reads overlapping exons in each gene were counted using featureCounts^75^, and these counts were used an input into DESeq2^76^. DeSeq2 function counts(normalized=TRUE) was used to calculate normalized counts with a regularized log transformation. The DESeq() and results() functions were used to calculate gene expression differences between pairs of neuronal populations.

### Accession numbers

Raw data is available at GEO. To review GEO accession GSE113663: Go to https://www.ncbi.nlm.nih.gov/geo/query/acc.cgi?acc=GSE113663 Enter token kzwxsgecxvkblqv into the box

## Acknowledgements

We thank all members of the Li and Shohat-Ophir labs for fruitful discussion and technical support. We specially thank Prof. Eli Eisenberg, Prof. Erez Levanon and Dr. Ulrike Heberlein for insightful comments on the manuscript. We also thank Dr. Fred Davis and Dr. Andrew Lemire for technical insights regarding the INTACT protocol, and Dr. Avi Jacob for technical support with imaging. This work was supported by the Israel Science Foundation (384/14), the United states – Israel Binational Science Foundation (2015275), National Institutes of Health (R01 GM102484, R01 GM124215, and R01 MH115080), National Science Foundation Graduate Research Fellowship (no. DGE-114747 to A.L.S.), and NIH training grant NIH-NIGMS T32 GM007790 (to A.L.S.).

## Author contributions

A.S., A.L.S., G.S.O, and J.B.L. conceived of experiments. A.S. performed INTACT RNA purifications with input from G.L.H. and RNA-seq library preparation. O.Y prepared RNA-seq libraries, T.S. assisted with sequencing and RNA-seq data analysis. A.L.S. performed mmPCR experiments and editing data analysis with input from Q.L. Y.P. performed structural modeling. A.S. and A.L.S. wrote the manuscript with input from G.S.O. and J.B.L.

## SUPPLEMENTARY INFORMATION

### Supplementary Note

#### Detailed description of novel editing site discovery

We mapped the RNA-seq reads using STAR^1^, combined the replicates of each sample for increased coverage, and called variants using GATK^2^, filtering out spurious variants as previously reported^3^ (**Supplementary Figure 1A**). We further required potential sites to be present in at least two of three replicates for each neuronal population. After filtering variants following our previously developed protocols, we found a large number of C-to-T and G-to-A variants and a calculated false discovery rate of 26%. We believed that DNA sequence differences between the parental fly strains of the sequenced F1 progeny were responsible for the large number of variants not caused by A-to-I editing. Because ADAR proteins often edit multiple adenosines in a region^4^, particularly in *Drosophila*^5^, we required putative editing sites to be adjacent to variants of the same type. This approach was recently shown to aid in distinguishing editing events from SNPs^6^. We further filtered variants, requiring two or more of the same type of base conversion within 200 bases of each other, and we found that using this additional filtering step greatly reduced the total number of false positive editing sites called from these datasets with a minimal decrease in sensitivity (**Supplementary Figure 1B**). The vast majority (97.7%) of *de novo* sites filtered out with this step had low coverage across most populations and therefore would not have been considered in our comparative analysis.

After all filtering steps, individual populations had between 85% and 95% A-to-G or T-to-C variants, indicating low numbers of false positive events in all samples. After combining all variants identified in all neuronal populations, 88% were A-to-G or T-to-C. By comparing the number of C-to-T and G-to-A changes to the number of A-to-G and T-to-C changes, we calculated a false discovery rate of 5.6%.

To determine which sites were novel, we compared our *de novo* identified sites to known sites from^7-16^. When comparing editing levels between the previously known and newly identified novel sites in the populations in which they were identified, we found that the two groups had similar median editing levels (29% and 26% respectively), but the distribution of editing levels of known sites skewed higher than the novel sites (**Supplementary Figure 1C**). The known sites tended to have higher sequencing coverage than the novel sites, as the median coverage of known sites was 105 reads and the median coverage of novel sites was 40 reads (**Supplementary Figure 1D**).

Because our *de novo* editing site discovery pipeline included stringent filters to avoid false positives, we also measured editing levels at all previously discovered and characterized editing sites in flies, whether we identified them *de novo* or not. Doing so allowed us to compare editing at an additional 435 editing sites between neuronal populations. Of those 435 previously known sites, 25 were identified *de novo* but filtered out because they were not found in clusters. The remaining 410 were filtered out earlier in our pipeline for various other reasons including being close to splice sites, within homopolymeric regions, or overlapping simple repeats, but since they had already been identified and showed reproducible editing levels, we used them in our comparative analysis. To obtain high coverage and accurate editing levels at additional known editing sites that were not covered by RNA-seq, we used mmPCR-seq to amplify sites of interest, and this added coverage of an additional 365 sites that were not identified as high confidence *de novo* sites.

### Supplementary Figures

**Supplementary Figure 1.**
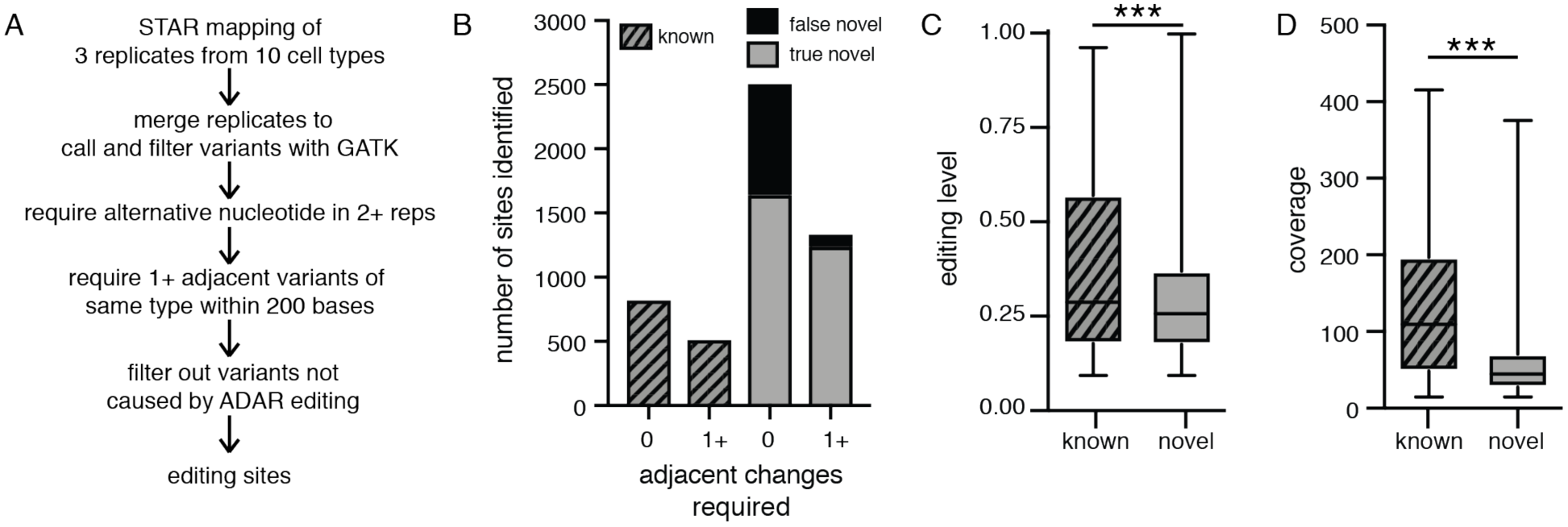
Pipeline to identify novel sites from RNA-seq. **(A)** Schematic of analysis pipeline for identifying novel editing sites from ten neuronal populations. **(B)** The total number of A-to-G or T-to-C variants identified *de novo* before and after requiring one or more adjacent variants of the same type. On the left are the number of previously known editing sites, on the right are the number of novel sites that are presumed to be true editing sites and the number of novel sites that are presumed false positives based on the calculated false discovery rate. **(C)** Box plot of editing levels at known and novel sites identified *de novo* in all populations. Whiskers show minimum to maximum values, with boxes representing 25th-75th percentile and median shown. *** p-value < 0.0001, two-tailed Mann-Whitney-U test. **(D)** Box plot of sequencing coverages at known and novel sites identified *de novo* in all population. *** p-value < 0.0001, two-tailed Mann-Whitney-U test.

**Supplementary Figure 2.**
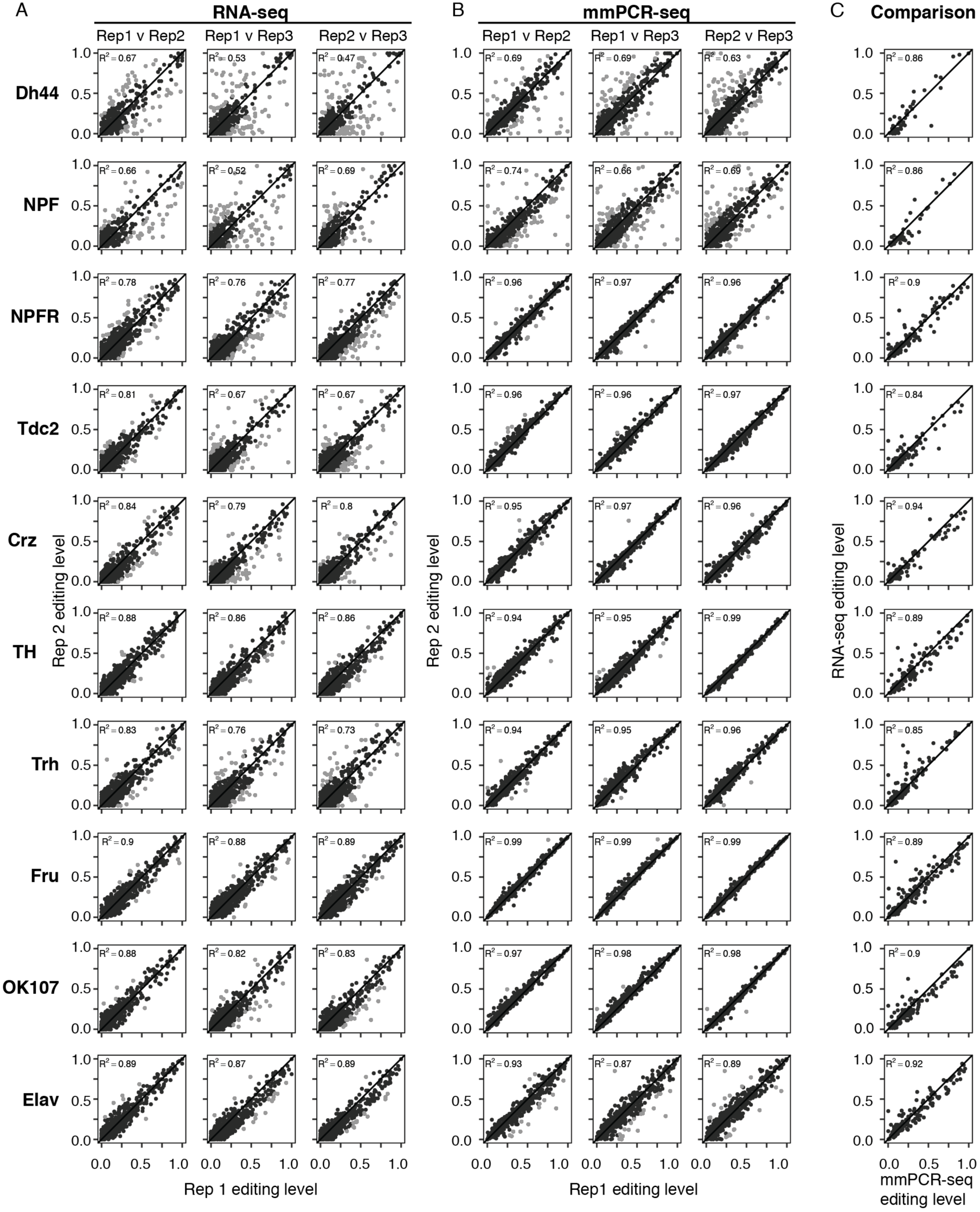
Editing levels measured from RNA-seq and mmPCR-seq are reproducible between replicates and consistent with each other. **(A-B)** Scatter plots of pairwise biological replicates from each of the cell types from RNA-seq **(A)** and mmPCR-seq **(B)**. Pearson’s correlations (R2) are shown. Sites where editing levels differed by >20% editing between replicates (light gray) were excluded from further analysis because these differences can be caused by technical artifacts, especially from populations with small numbers of neurons like Dh44 and NPF. **(C)** Scatter plot comparisons of the average editing levels between RNA-seq replicates and mmPCR-seq at the subset of sites covered in both in each cell type.

**Supplementary Figure 3.**
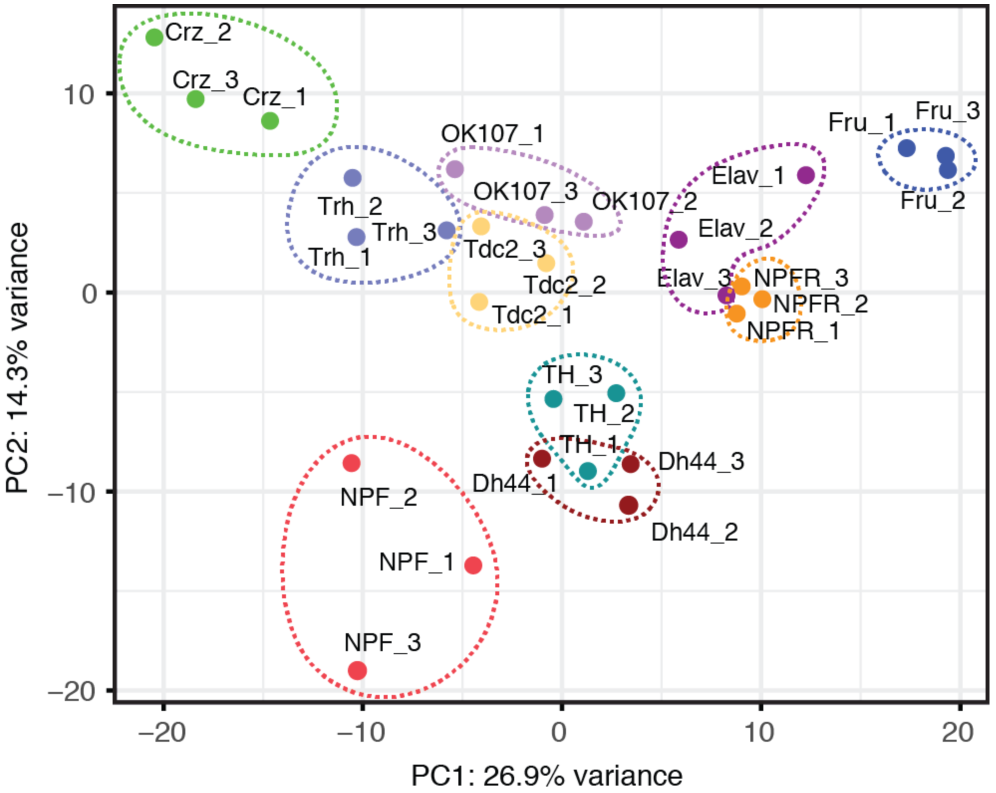
Principal component analysis of editing levels across populations. Principal component analysis of editing levels as measured by either mmPCR-seq or RNA-seq in all replicates at sites that are reproducible between replicates and covered in all samples.

**Supplementary Figure 4.**
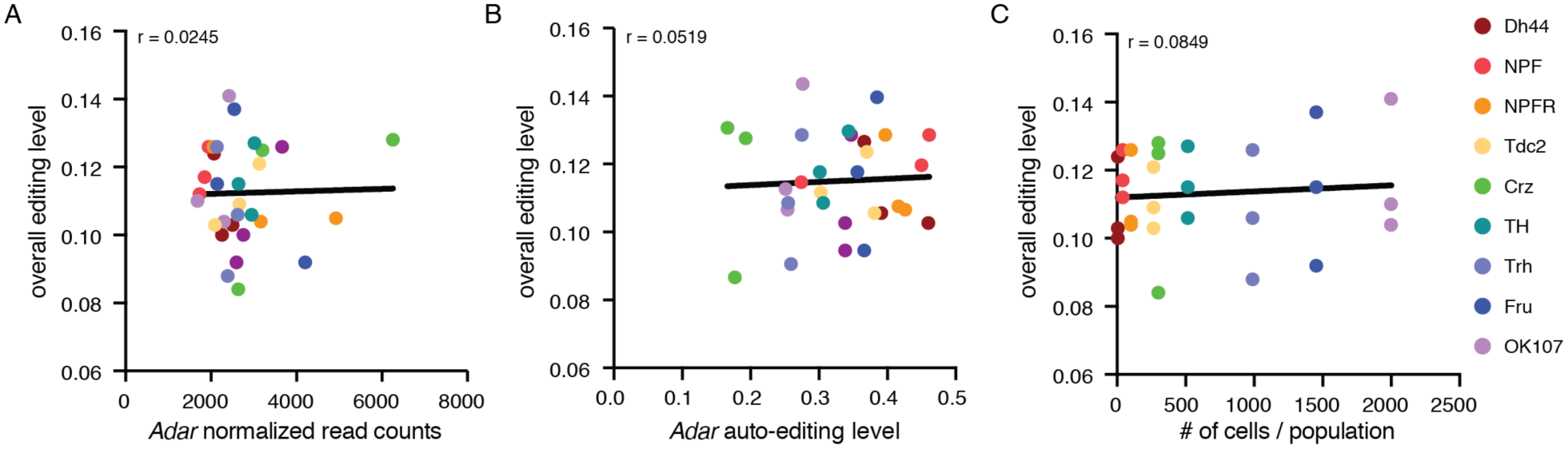
Overall editing levels do not correlate with *Adar* expression, *Adar* auto-editing, or the number of cells per population. **(A)** Scatter plot of overall editing level versus to *Adar* normalized read counts for each replicate of RNA-seq. Overall editing levels were calculated as the number of G reads at all known editing sites over the total number of reads at all known editing sites for each replicate. **(B)** Scatter plot of overall editing level versus *Adar* auto-editing levels for each replicate of RNA-seq. **(C)** Scatter plot of overall editing level versus the number of cells in each neuronal population for each replicate of RNA-seq. Pearson correlations and linear regression lines calculated in GraphPad PRISM 7 are shown.

**Supplementary Figure 5.**
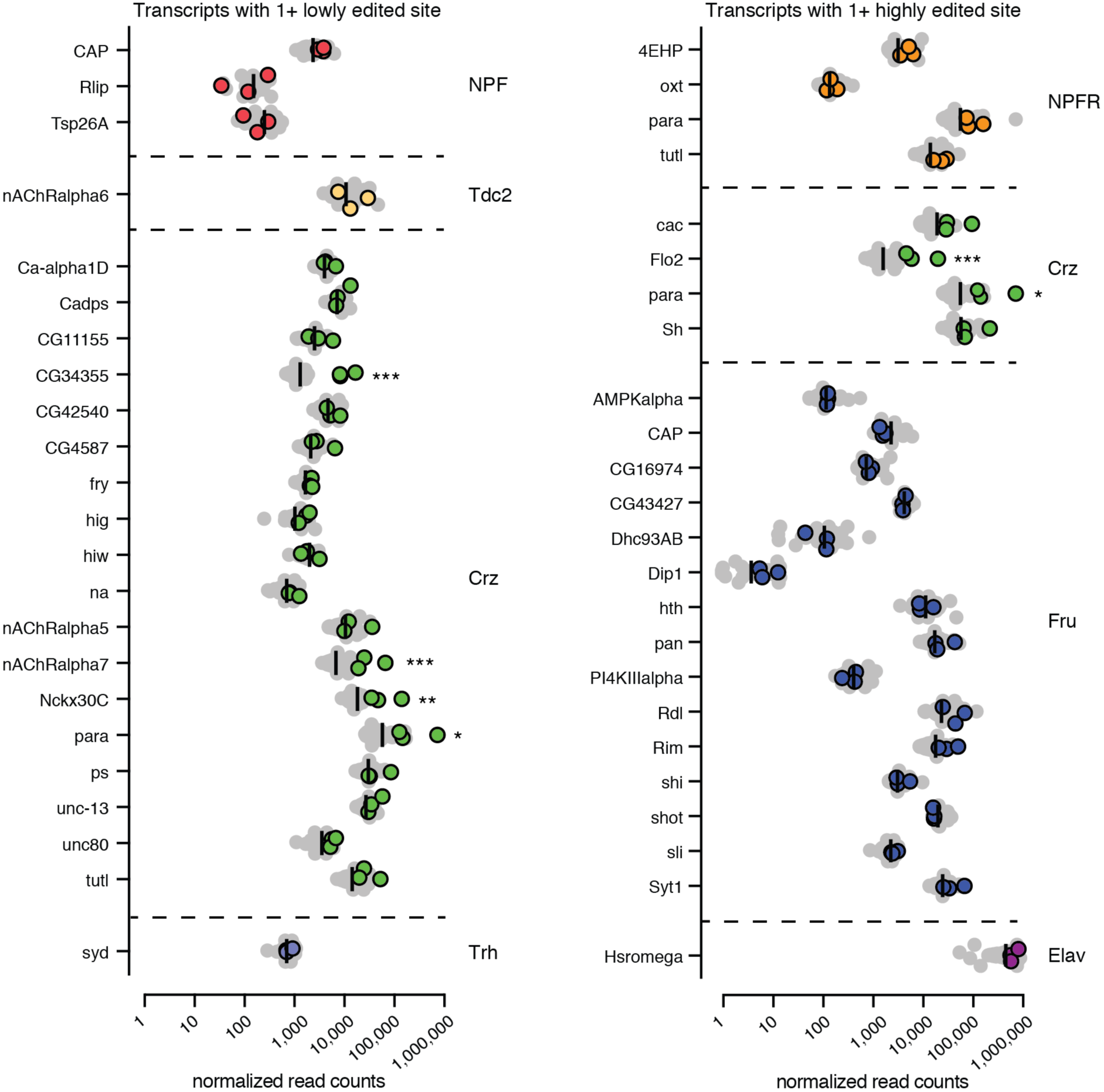
Expression of transcripts with population-specific editing. Normalized counts of transcripts that have population-specific editing, grouped by the population in which editing is unique (listed on right). Colored dots are three replicates from the population with specific editing, gray are replicates from all other populations, black lines are median counts of all populations. Transcripts with lowly edited sites are on left, and transcripts with highly edited sites on right. X-axis is log10 scale. Significant expression differences were determined through pairwise comparisons between expression in highlighted population versus all other populations using DESeq2. P-values were calculated using Wald tests. * p-value < 0.05 in all pairwise comparisons, ** p-value < 0.01 in all pairwise comparisons, *** p-value < 0.001 in all pairwise comparisons.

**Supplementary Figure 6.**
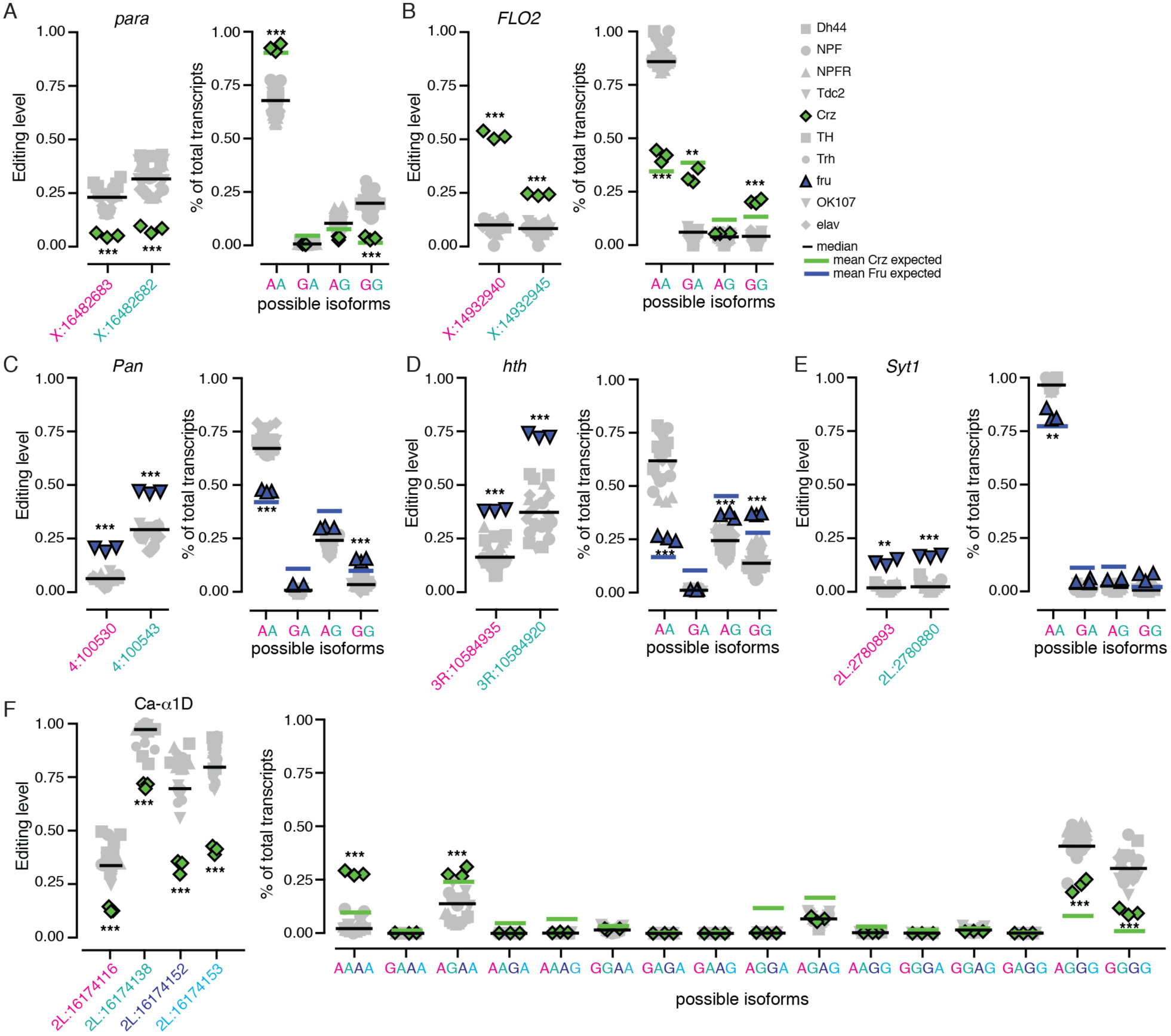
Co-regulation of editing sites. **(A-B)** Editing levels (left) and isoform usage (right) in clusters of editing sites that are differentially regulated in Crz neurons in *para* and *FLO2*. **(C-E)** Editing levels (left) and isoform usage (right) in clusters of editing sites that are differentially regulated in Fru neurons in *Pan*, *hth*, and *Syt1* transcripts. **(F)** Editing levels (left) and isoform usage (right) in cluster of four editing sites that are differentially regulated in Crz neurons in *Ca-alpha1D*. Crz and Fru editing and isoform usage are shown in green and blue respectively, with other populations in gray. Black bars are median editing level of all populations, green and blue bars represent mean expected usage of each isoform in Crz and Fru respectively based on editing levels of the clustered sites. ** p-value < 0.01, *** p-value < 0.001 from Welch’s t-test. Editing site locations are noted as chromosome: position.

### Supplementary Tables

**Supplementary Table 1. Coverage and editing levels of editing sites discovered de novo from RNA-seq.** (see Supplementary_Table1.xlxs)

**Supplementary Table 2. A and G counts at editing sites and Fisher’s exact test p-values from pairwise comparisons between populations.** (see Supplementary_Table2.xlxs)

**Supplementary Table 3. Editing levels, z-scores and p-values from Welch’s T-test for each replicate of all populations.** (see Supplementary_Table3.xlxs)

**Supplementary Table 4. GO Term enrichment for transcripts that contain Crz-specific editing events.** (see Supplementary_Table4.xlxs)

**Supplementary Table 5. Normalized read counts and log2 fold change calculations and p-values between populations for marker genes, Adar, and transcripts with population-specific editing.** (see Supplementary_Table5.xlxs)

**Supplementary Table 6. Observed and expected isoform usage of clustered editing sites with p-values from T-tests.** (see Supplementary_Table6.xlxs)

**Supplementary Table 7.** RefSeq and Uniprot ID numbers for transcripts with editing differences in specific neuronal populations.

| Gene Name | RefSeq Transcript ID | Uniprot ID |
| --- | --- | --- |
| *Ca-alpha1D* | NM_134429 | Q24270 |
| *cacophony* | NM_001258710 | P91645-1 |
| *CG4587* | NM_001201888 | A8DZ06 |
| *na* | NM_001103511 | A8JUW5 |
| *unc80* | NM_143320 | Q9VB11 |
| *nAChRalpha6* | NM_205953 | Q7KTF8 |
| *nAChRalpha7* | NM_143320 | Q9VWI9 |
| *nAChRalpha5* | NM_001259098 | Q7KT97 |
| *paralytic* | NM_078647 | P35500-1 |
| *Shaker* | NM_001272736 | P08510-1 |

